# A two-stage approach for the spatio-temporal analysis of high-throughput phenotyping data

**DOI:** 10.1101/2021.08.10.455613

**Authors:** Diana M. Pérez-Valencia, María Xosé Rodríguez-Álvarez, Martin P. Boer, Lukas Kronenberg, Andreas Hund, Llorenç Cabrera-Bosquet, Emilie J. Millet, Fred A. van Eeuwijk

## Abstract

High throughput phenotyping (HTP) platforms and devices are increasingly used for the characterization of growth and developmental processes for large sets of plant genotypes. Such HTP data require challenging statistical analyses in which longitudinal genetic signals need to be estimated against a background of spatio-temporal noise processes. We propose a two-stage approach for the analysis of such longitudinal HTP data. In a first stage, we correct for design features and spatial trends per time point. In a second stage, we focus on the longitudinal modelling of the spatially corrected data, thereby taking advantage of shared longitudinal features between genotypes and plants within genotypes. We propose a flexible hierarchical three-level P-spline growth curve model, with plants/plots nested in genotypes, and genotypes nested in populations. For selection of genotypes in a plant breeding context, we show how to extract new phenotypes, like growth rates, from the estimated genotypic growth curves and their first-order derivatives. We illustrate our approach on HTP data from the PhenoArch greenhouse platform at INRAE Montpellier and the outdoor Field Phenotyping platform at ETH Zürich.

## Introduction

In recent years, the frequency of use of high-throughput phenotyping (HTP) techniques and platforms has strongly increased in plant genetics and physiology. HTP data provide quick, precise, non-destructive and cost-effective information on phenotypic traits with high spatial and temporal resolution^1^. Designed HTP experiments, either indoors or in a field, usually consist of experimental units (e.g., single plants in pots or plots) that are combined with a wide range of sensing equipment for the (almost) continuous monitoring of plant/plot phenotypic traits for large sets of genotypes. Treatments applied in HTP research designs may comprise not only different genotypes but also different management practices. High dimensional HTP data as derived from multiple sensors (e.g., images, point clouds, hyperspectral data) are typically filtered, condensed, integrated and summarised into features. Combinations of one or more features are used to approximate simple biological traits that are on the one hand still close to the data, and on the other hand are relatively far from the target traits of commercial interest (most often yield and quality parameters). Examples of such low-level traits are plant height, canopy cover, leaf area index, ear and tiller counts, canopy temperature or indices related to water or chlorophyll content. Thus, researchers and plant breeders have now access to large and detailed datasets, in the form of (long) time-series, enabling to track multiple low-level traits from, e.g., seed emergence to physiological maturity. The challenge in this setting is how to efficiently and adequately exploit the diversity and complexity of HTP data to (a) extract relevant information on several aspects of plant growth and development, (b) increase the biological understanding of plant systems; and (c) help and support the decision-making process in plant breeding programs^2^. On top of that, the vast amount of data typically produced by HTP platforms require not only appropriate but also rapid processing techniques.

Regardless of the fact that complex traits (like yield) are measured only once (as traditionally) or low-level phenotypic traits are measured several times (as in HTP), it is known that their expression is spatially affected by environmental factors such as soil heterogeneity in field experiments and temperature and light gradients in the greenhouse. This makes it necessary to correct for these nuisance factors when analysing agricultural experiments^2, 3^. For “traditional” agricultural experiments, modelling spatial variation through spatially correlated stochastic components or smoothing techniques has become ubiquitous^4–11^. Yet, data from HTP experiments present, on top of spatial variation, a time dimension which needs to be incorporated and adequately modelled in the analyses. Although interest may primarily focus on estimating (and further processing and analysing) the evolution over time of the genetic effect on a given phenotype, it will be essential to correct for environmental nuisance effects not only in a spatial but now also in a temporal dimension. Thus, from the statistical point of view, HTP data ask for spatio-temporal modelling approaches. To the best of our knowledge, there are few proposals in the HTP literature that tackle the problem from both perspectives, temporal and spatial. References for stage-wise analysis of HTP data include the work by van Eeuwijk *et al*. (2019)^2^, where the authors propose to first estimate time-series of spatially adjusted genotypic means from low-level phenotypic traits, that are then subject to temporal dynamic modelling. A similar approach is followed by Kar *et al*. (2020)^12^. In the above-cited references, the information that is carried to the temporal analysis is not at the level of the experimental unit but at the genotype level. A different stage-wise approach is described in Roth *et al*. (2021), where the temporal analysis is performed first, followed by the spatial correction. Approaches that model simultaneously the spatial and temporal genetic and non-genetic variation in one-stage include the works by Verbyla *et al*. (2021)^13^ and Pérez *et al*. (2020)^14^. While promising, both proposals report high computational complexity and scalability problems.

In this work we propose a two-stage approach that allows keeping the data resolution for the second stage at the experimental unit (plant/pot/plot) and is computationally efficient. The first stage focuses on correcting the low-level phenotypic data for experimental design factors and spatial variation, while the second stage aims at estimating the evolution over time of the genetic signal. In a similar fashion to van Eeuwijk *et al*. (2019)^2^, in the first stage we fit a spatial model to the phenotypic data separately for each measurement time. The phenotypic data are subsequently corrected by only considering the (estimated) sources of variation which are of interest plus the residual component (i.e., the measurement error). The purpose of this stage and subsequent correction is two-fold: (1) to remove nuisance spatial variation from the phenotypic data; and (2) to keep the data resolution for the second stage at the level of the experimental unit (through the incorporation in the correction of the residual component). This is one of the main differences to the proposals described in van Eeuwijk *et al*. (2019)^2^ and Kar *et al*. (2020)^12^, and it is routinely applied for data derived from the field phenotyping platform of ETH Zurich^15–17^. Since analyses are performed separately for each measurement time, our modelling strategy implicitly permits the spatial variation to differ among measurement times, i.e., it allows correcting for both the spatial and temporal evolution of environmental variables and experimental design factors. For this first stage of spatial correction, different proposals are available in the literature^4–11^. The separable autoregressive model^5, 6^ has become the standard modelling strategy, but the recently proposed SpATS (Spatial Analysis of Field Trials with Splines) approach^11^ has proved to be a powerful alternative. Indeed, comparisons of these two models have shown that SpATS is a very competitive approach, with the advantage that model selection is greatly simplified^18, 19^. In addition, an `R`-package (also called `SpATS`) is freely available on CRAN (https://CRAN.R-project.org/package=SpATS), and the recently `statgenHTP`^20^ `R`-package (available on https://CRAN.R-project.org/package=statgenHTP) allows for an easy fitting of SpATS models for different (and possibly large) measurement times. For all these reasons, we choose SpATS for this first stage.

The second stage of our proposal focuses on modelling the genetic signal as a function of time for the corrected phenotype obtained in the first stage. Data for this stage consist of time-series of spatially corrected phenotypic trait measurements per experimental unit. In other words, we have a sample of growth curves per plant/pot/plot that can be grouped by, at least, one treatment factor, the genotype^21^. Thus, generally speaking, this stage requires hierarchical or multilevel growth curve modelling approaches. A traditional data-driven approach to the analysis of growth curves is to use parametric (non-linear) models, being the logistic function one of the most commonly used^22^. However, while growth processes theoretically follow a clearly defined pattern that may be modelled using a parametric function, the observed dynamics may deviate considerably due to, for example, temporal changes in environmental conditions (e.g. cold spells) or the application of treatments (e.g. irrigation events). More flexible models that overcome the limitations of parametric specifications have been proposed in the literature. Examples include the use of smoothing or penalised splines (P-splines^23^) as well as functional principal components analysis (FPCA)^21, 24, 25^. For the particular case of data from HTP platforms, their temporal analysis by means of splines or FPCA has been discussed in Xu *et al*.^26–28^, Wang *et al*.^29^ and Brien *et al*.^30^. In this paper, we follow the same modelling philosophy as for the first stage and propose the use of P-spline hierarchical curve data models along the lines of the work by Durban *et al*. (2005)^24^ and Greven and Scheipl (2017)^31^. Our approach presents several advantages. The connection between P-splines and linear mixed model offers a rich framework for estimation and inference^32, 33^. Moreover, we can easily obtain the derivatives of the estimated curves at different levels of the hierarchy (e.g., genotype or experimental unit) which can provide important insights on genotype performance. Finally, from the estimated curves and their derivatives, different time-dependent and -independent characteristics (intermediate traits) can be easily extracted. Intermediate traits capture the dynamics of low-level traits and are closer to the target traits of plant breeders. Examples include the area under estimated growth curves, the timing of key stages (e.g., the timing at which final height is reached), or quantities at defined time points or periods (e.g., final height, final canopy cover or maximum speed rates)^34^. They can serve as inputs to subsequent statistical analyses that aim at modelling genotype-by-environment interactions in biologically complex traits, like yield, in terms of underlying component traits.

## Methods

We consider that, in the HTP experiment, genotypes have been allocated to the experimental units following an experimental design. For simplicity, in what follows we assume that only genotypes have been tested (i.e., we have a single factor treatment structure). However, we allow for population structure, modelled as different families, panels or populations of genotypes. Thus, the data present a three-level nested hierarchical structure, with plants/plots nested in genotypes, and genotypes nested in populations (see Supplementary Fig. S1 online). For clarity and simplicity, hereafter we refer to the experimental units of the experimental design as plants. The experimental unit is most commonly a plant for an indoor experiment and a plot containing several plants for a field experiment.

### Data Structure

Let *y*_*pgi*_(*t*) denote the (observed) low-level phenotypic trait of interest for the *i*-th plant (*i* = 1, …, *m*_*pg*_) of the *g*-th genotype (*g* = 1, …, ℓ_*p*_) in the *p*-th population (*p* = 1, …, *k*) at time *t* ∈ {*t*_1_, …, *t*_*n*_}. Let 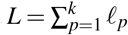 denote the total number of genotypes and 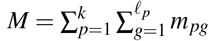 the total number of plants. We consider that plants can be mapped to a coordinate system defined in terms of *r* rows and *c* columns, and denote as *u*_*pgi*_ and *v*_*pgi*_ the row and column position respectively (*u*_*pgi*_ ∈ *{*1, …, *r*} and *v*_*pgi*_ ∈{1, …, *c*}). In the first stage, the correction is made per time point for the whole experiment. As such, note that we assume that all plants in the experiment are measured at the same times, (*t*_1_, …, *t*_*n*_). That is a simplification, as platform data is typically acquired within the order of minutes to hours. However, we presume that the factors that may affect the platform measurements within that period can be accounted for (and captured) by the experimental design (e.g., blocking structure). The assumption of the same measuring times, however, does not preclude the presence of incomplete data. The methods can handle missing values at both the plant and genotypic level (i.e., with plants or even genotypes not measured for some times).

### First stage: Environmental Correction using SpATS

In the first stage, a SpATS model is fitted to the phenotypic data separately for each measurement time *t* ∈{*t*_1_, …, *t*_*n*_}. A SpATS model is a linear mixed model where large-scale and small-scale spatial dependence (or spatial trend) is explicitly modelled by a two-dimensional smooth surface defined over the row and column positions of the plants in the experiments, *f*_*t*_(*u, v*). This smooth function is constructed with tensor-product P-splines^23, 35^, i.e., through the combination of the tensor-product of marginal B-spline bases and an anisotropic discrete penalty on the coefficients. In its more general specification, and considering genotypes as random, a SpATS model has the following form (for more details, see Rodríguez-Álvarez *et al*.^11^)

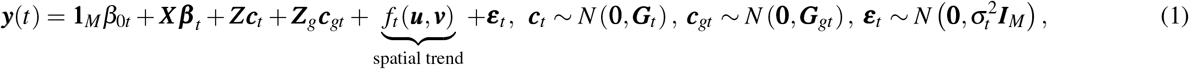

where 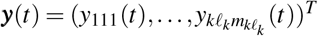 is the low-level phenotypic trait at time *t*, ***X*** and ***Z*** represent column-partitioned matrices, associated respectively with fixed and random components, as for instance, row, column, replicate and/or (incomplete) block effects, and ***Z***_*g*_ is the design matrix assigning plants to genotypes. Further, we decompose the fixed effects component in two terms

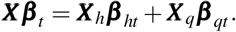

The first term, ***X***_*h*_***β***_*ht*_, corresponds to the factors/covariates whose effects we are interested in modelling in the second stage, whereas ***X***_*q*_***β***_*qt*_ corresponds to the factors/covariates whose effects we are interested in removing (e.g., those associated with experimental design factors). As said before, for simplicity, we consider that ***X***_*h*_***β***_*ht*_ only contains information regarding the family/population to which the genotypes belong to (if any). Concerning ***X***_*q*_***β***_*qt*_, we assume that it is associated with *R* experimental design factors (categorical covariates), and thus, ***X***_*q*_ = [***X***_*q*1_ | … | ***X***_*qR*_] and 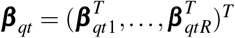. The length of ***β***_*qtr*_ (and therefore the number of columns in the associated design matrix ***X***_*qr*_) (*r* = 1, …, *R*) corresponds to the number of different categories, say *c*_*r*_, of the *r*-th experimental design factor minus one (as the intercept is included in the model).

Once the SpATS model in equation (1) is fitted, the phenotype of interest at time *t*, ***y***(*t*), is corrected by only considering the (estimated) sources of variation that are of interest, plus the residual component, i.e., the corrected phenotype, denoted as 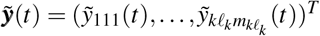, is obtained as follows

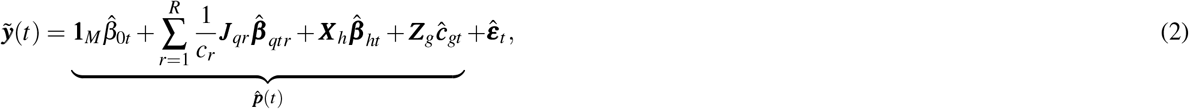

where ***J***_*qr*_ are matrices of ones of appropriate dimensions (i.e., *M* × (*c*_*r*_ −1)). The correction is performed following the procedure for obtaining predictions (e.g., adjusted means) in linear mixed models described in Welham *et al*. (2004)^36^. In that paper, the authors propose a partition of the explanatory variables (see equation (1)) in three groups: (1) those for which predictions are required (i.e., population, 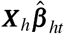, and genotypic, ***Z***_*g*_***ĉ***_*gt*_, effects), (2) those to be averaged over (i.e., experimental design factors effects, 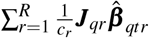); and, (3) those to be ignored (i.e., spatial trends, 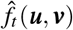, and other random effects, ***Zĉ***_*t*_).

As will be seen in the next section, in the second stage of our proposal we model 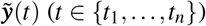. Thus, it is worth emphasising that, in the way it is constructed, 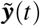 (see equation (2)) only contains information about genetic populations and genotypes, as well as unexplained plant-to-plant variation (measurement error). In other words, for the second stage the predicted values for the genetic populations and the genotypes, 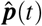, as well as the unexplained plant-to-plant variation, 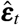, are maintained as the “new” (spatially corrected) experimental unit values, while the spatial trends and other blocking factors to control for spatial variability are omitted. Also, note that the “observations” that enter the second stage, 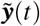, are not observed but estimated/predicted. Thus, we propose to propagate the uncertainty from the first stage to the second stage through the inclusion of weights, in a similar way to the weighted stage-wise analysis of multi-environment trials^37^. In particular, weights are obtained from the inverse of the variance-covariance (vcov) matrix for the predictions (plus the residual variance), i.e., 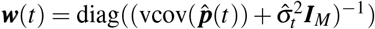.

### Second stage: P-spline Hierarchical Curve Data Model

The aim of the second stage is to model the spatially corrected phenotype obtained in the first stage. We can re-organise the data for this stage in such a way that they can be seen as a sample of plant growth curves, 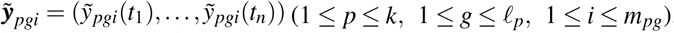, with a nested hierarchical structure, where plants are nested in genotypes and genotypes are nested in populations. We propose to model this sample of curves by considering an additive decomposition of the phenotypic variation over time and use a three-level nested hierarchical growth model for this purpose^21^

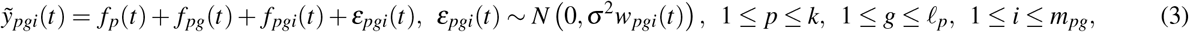

where *f*_*p*_ is the growth/change over time of the (spatially corrected) phenotype for the *p*-th population (i.e., the *p*-th population mean function), *f*_*pg*_ is the genotype-specific growth deviation from *f*_*p*_ for the *g*-th genotype, and *f*_*pgi*_ is the plant-specific growth deviation from *f*_*pg*_ for the *i*-th plant. The additive modelling approach implies that *f*_*p*_ + *f*_*pg*_ can be interpreted as the growth over time of the (spatially corrected) phenotype for the *g*-the genotype in the *p*-th population. Thus, on top of genotype-specific deviations from their overall population mean, we also obtain genotype-specific growth curves. Finally, *w*_*pgi*_ are the weights obtained in the first stage.

We use P-splines for hierarchical curve data^24, 31^ to estimate the model in equation (3). In this framework, each function in equation (3) is approximated by a linear combination of cubic B-spline basis functions, and its smoothness is controlled by a penalty on the differences of the B-spline coefficients (we use differences of order 2). The influence of the penalty is determined by a smoothing parameter. One of the attractive features of using B-splines to represent the functions in equation (3) is that their derivatives can be easily obtained^38^. Furthermore, we adopt the connection between P-splines and linear mixed models^32, 33^, where each function is treated as a sum of fixed (linear) and random (non-linear) components, and the smoothing parameter is replaced by a ratio of variances components. For clarity we omit here the more technical details, but it can be shown that, under this setting, each function in equation (3) is expressed as follows

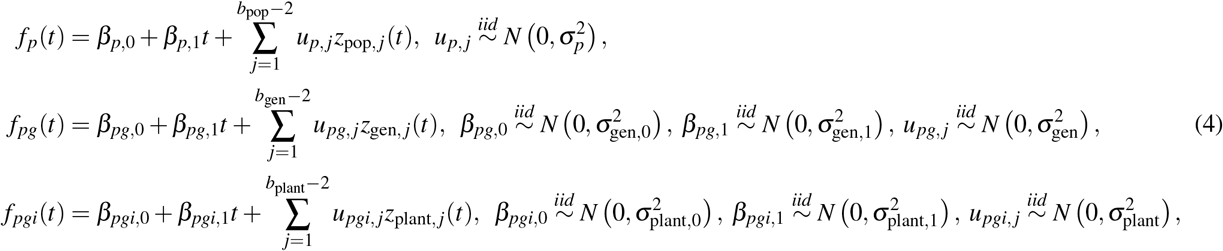

1 ≤ *p* ≤ *k*, 1 ≤ *g* ≤ ℓ_*p*_, 1 ≤ *I* ≤ *m*_*pg*_, where {*z*_pop, *j*_(·) : 1 ≤ *j* ≤ *b*_pop_ – 2}, {*z*_gen, *j*_(·) : 1 ≤ *j* ≤ *b*_gen_ – 2} and {*z*_plant, *j*_(·) : 1 ≤ *j* ≤ *b*_plant_ – 2} are sets of basis functions obtained from the connection between P-splines and linear mixed models^32^. In contrast to standard P-spline mixed models, here the linear component (intercept and slope) associated with *f*_*pg*_ (genotypic deviations) and *f*_*pgi*_ (plant deviations) is modelled with random rather than fixed effects. Also, for notational simplicity, we assume the same genetic variation across populations. However, this assumption can be easily relaxed by considering different values of 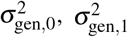 and 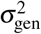 per population. A similar approach can be followed to allow for the plant-to-plant variation (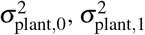 and 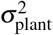) to vary across genotypes. These generalisations might be worth exploring if there are sufficient number of genotypes per population and plants per genotype, respectively.

Before proceeding, we comment on the selection of the number of B-spline basis functions used to approximate *f*_*p*_, *f*_*pg*_ and *f*_*pgi*_ (i.e., *b*_pop_, *b*_gen_ and *b*_plant_, respectively). In P-splines, it is recommended to choose a large number of bases to provide enough flexibility; the role of the penalty is to avoid over fitting^23^. In our setting the number of functions in the complete model equals *k* + *L* + *M* (populations + genotypes + plants), and, hence, the number of regression coefficients (either fixed or random) to be estimated is *k* × *b*_pop_ + *L* × *b*_gen_ + *M* × *b*_plant_. This value can be very large, with the number of plants, *M*, and associated basis dimension, *b*_plant_, playing the major role: the dataset may contain thousands of plants. Thus, to reduce the computational burden, one could be tempted to use different basis dimensions for *f*_*p*_, *f*_*pg*_ and *f*_*pgi*_, and to be less generous with *f*_*pgi*_. However, this is not a good strategy. Simulation studies, as well as preparatory data analyses, have shown that results may be sensitive (and in some cases unreliable) to using different bases dimensions. We therefore recommend choosing the same value for *b*_pop_, *b*_gen_ and *b*_plant_, while keeping the number of coefficients at a reasonable level (i.e., a trade-off between flexibility and dimensionality).

We now present the model in equation (3) based on the specification in equation (4) in matrix notation. Let’s first order the data by population, genotype, plant, and time (in that order), i.e., 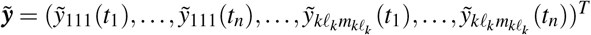. Thus, in a compact way, the three-level nested hierarchical growth model can be expressed as

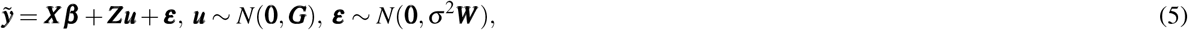

where

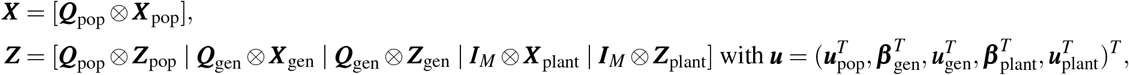

and

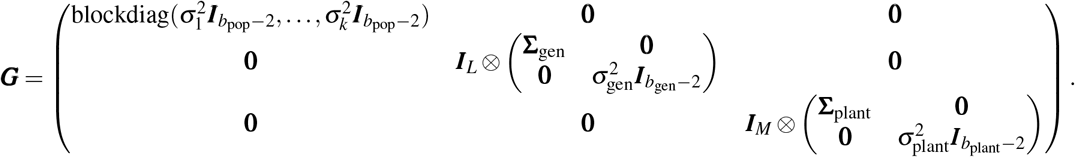

The specific form of the matrices and vectors involved in equation (5) is given in Section “Three-level Nested Hierarchical Growth Model: Mixed Model Formulation”. Note that model in equation (5) is a standard linear mixed model, and, thus, estimation can be carried out with any mixed-model software, such as the `R`-packages `ASReml-R`^39^, `nlme`^40^ and `lme4`^41^, or the `PROC MIXED`^42^ procedure in SAS^®^. Also, we can use the `R`-package `mgcv`^43^ for that purpose. However, high-throughput phenotypic data are usually characterised by a large number of observations, which, together with the number of regression coefficients in equation (5), might make estimation with the above-mentioned software computationally expensive. Thus, we have implemented in the `R` language^44^ our own code (provided along with the paper), which resorts to the recently proposed SOP (Separation of Overlapping Penalties) method^45^. Empirical best linear unbiased estimates (BLUE) and predictors (BLUP) are obtained by the solution of Henderson’s mixed model equations^46^, and variance components by means of restricted maximum likelihood (REML)^47^. Construction of (approximate) confidence intervals for the estimated curves and their derivatives is based on the prediction error variance^36^. To speed up computation, we take advantage of the array structure of the data through Generalised Linear Array Models (GLAM)^48^ and of the sparse structure of the matrices involved in the model. It is to note that the model in equation (5) presents a standard variance-covariance matrix for the random effects (i.e., it is linear in the variance parameters) and thus, the SOP method reduces to the estimating algorithm described in Harville (1977)^49^.

### Applications

We illustrate the potential of our approach with data from two experiments from two different HTP platforms: 1) the PhenoArch platform (INRAE Montpellier)^50^ (greenhouse, Figure 1(a)) and, 2) the FIP (FIeld Phenotyping) platform (ETH Zürich)^51^ (field, Figure 1(b)). The PhenoArch platform data are used to describe our approach and illustrate the outcomes at each step. The FIP platform data are further analysed to discuss the extraction of important (genotype-specific) attributes from the estimated genotype-specific growth curves and their first-order derivative, as well as their possible use in the decision-making process^2, 52, 53^.

**Figure 1.**
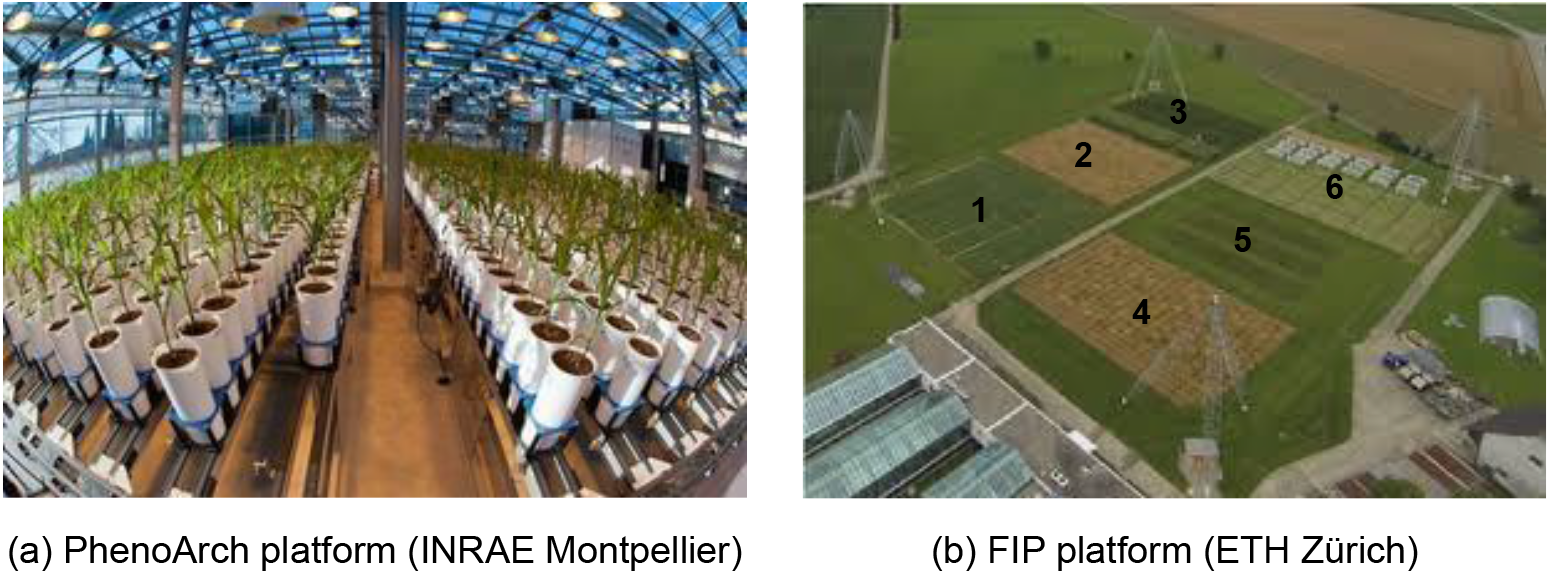
Overview of the two high-throughput phenotyping platforms.

#### PhenoArch platform (INRAE Montpellier)

The PhenoArch platform hosted at M3P, Montpellier Plant Phenotyping Platforms (https://www6.montpellier.inra.fr/lepse/M3P) is composed of a conveyor belt structure of 28 lanes carrying 60 carts with one pot each (i.e. 1680 pots on a rectangular grid of *r* = 60 rows and *c* = 28 columns (see Figure 1(a)), plus a conveyor belt system that feeds the imaging or the watering units. Pots are daily moved to be imaged and/or watered. They are then moved back to the same positions and orientation, so that the plant position with respect to its neighbours is conserved throughout the experiment. The data analysed here correspond to an experiment including two different panels of commercial maize hybrids representative of breeding history in Europe and US during the last 60 years. This material covers a wide range of plant architecture, growth and development. A total of 60 genotypes were tested in Panel 1 and 30 genotypes were tested in Panel 2, and each genotype was replicated between 4 (Panel 2) and 14 (Panel 1) times. All genotypes were tested under two levels of soil water content: (1) mild water deficit (WD, soil water potential of − 0.5 MPa) and (2) retention capacity (WW, soil water potential of 0.05 MPa). The experiment was carried out in 2017 between April 13th and May 15th, which correspond to 103 and 135 days since January 1st (hereafter referred as DOY, Day of the Year), respectively. Red-green-blue (2056 × 2454) images taken from 13 views (12 side views from 30° rotational difference and one top view) were captured daily for each plant. Plant pixels from each image were segmented from those of the background and used for estimating the whole plant leaf area (among other features) as described in Brichet *et al*. (2017)^54^. The dataset consists of *n* = 32 leaf area measurements on *M* = 1656 plants (1656 × 32 = 52992 observations, including missing data). For a proper analysis of this dataset, panels and treatments should be modelled as crossed effects. However, for the sake of simplicity, for our analyses we combined panels and water regimes, such that we considered *k* = 4 “populations” (Panel 1 - WD, Panel 1 - WW, Panel 2 - WD and Panel 3 - WW) and *L* = 180 “genotypes” (60 genotypes in Panel 1 and WW treatment, 60 genotypes in Panel 1 and WD treatment, 30 genotypes in Panel 2 and WW treatment and 30 genotypes in Panel 1 and WD treatment).

#### First stage results: PhenoArch platform

A SpATS model was fitted for each individual measurement time point of the leaf area data. In addition to the spatial trend and the random genotypic effects (see equation (1)), the model included the population (panel by water regime combination) as fixed effect, and the row and column positions as random effects. A different genetic variance for each of the four populations (panel by water regime combination) was considered. Regarding the spatial trend (i.e., the tensor-product P-spline), B-spline bases of dimension 60 and 28 were chosen for the row and column positions, respectively, and nested bases, with half the dimension, were used^55^. The spatially corrected leaf area included the estimated population and genotypic effects, as well as the residual (see equation (2)). The comparison of the spatially corrected leaf area when modelling genotypes as fixed (as usually in stage-wise analyses) or random (as done here) effects shows essentially identical results (Supplementary Fig. S3 online): shrinkage of the genotypic BLUPs is counteracted by the inclusion of the residual component into the correction in equation (2). Results show that, as expected, the spatial pattern observed in the raw data disappears in the spatially corrected phenotype, as illustrated with the leaf area at four different measurement times (Figure 2). Moreover, the correction reduced the variability among plants (i.e., replicates) of the same genotype and water treatment combination (Figure 3).

**Figure 2.**
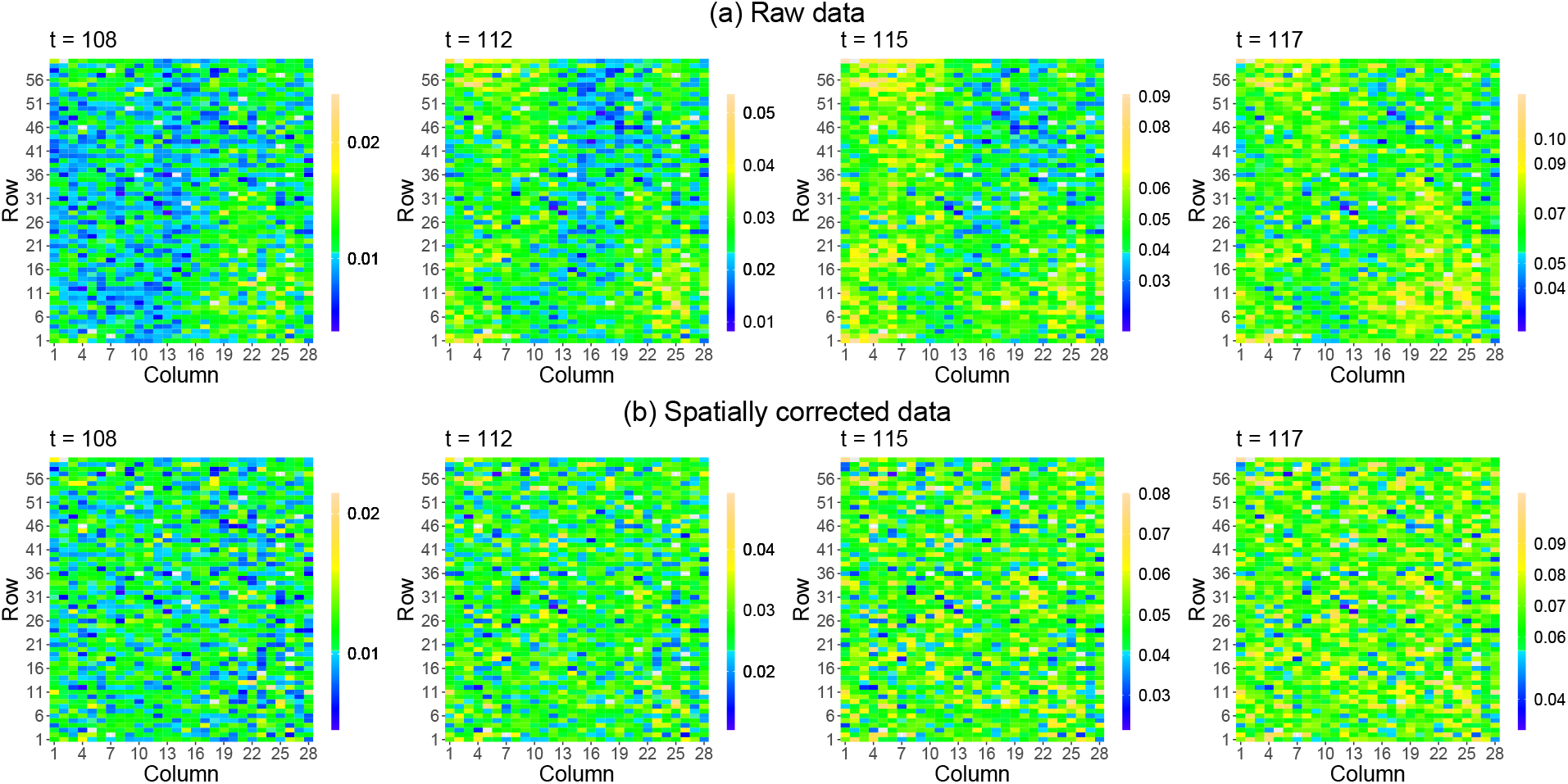
Results of the first stage of analysis for the PhenoArch platform: Spatial distribution of the (a) raw and (b) spatially corrected leaf area at four different measurements times (*t* = 109, 113, 116, 118 DOY). The white areas denote missing data. The colour scale is different for each time point.

**Figure 3.**
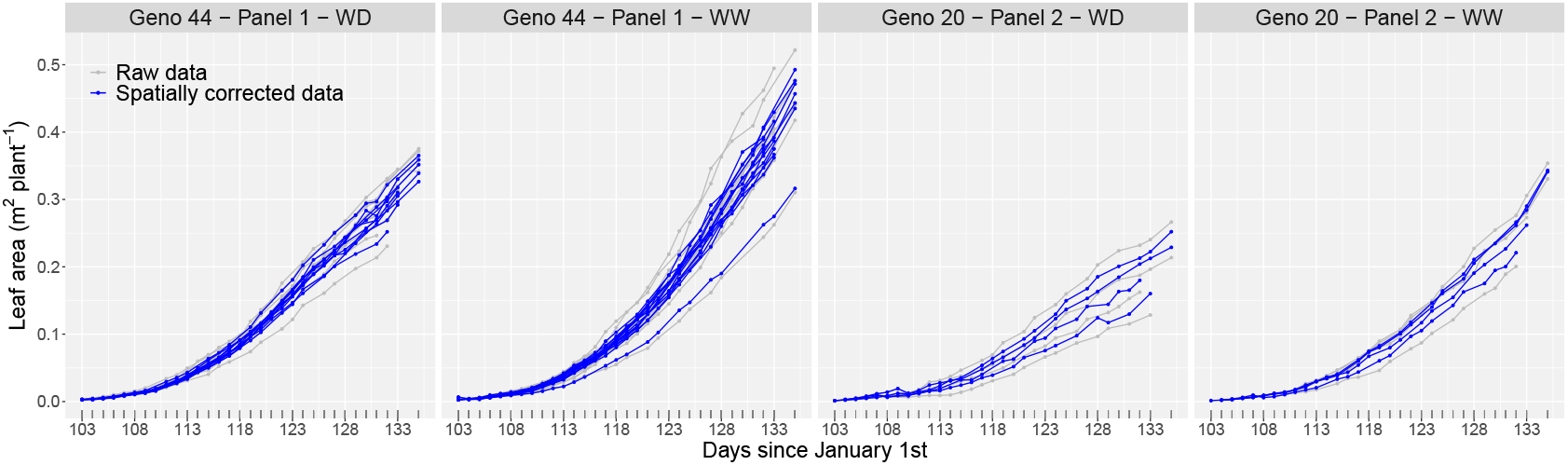
Results of the first stage of analysis for the PhenoArch platform: Evolution over time of the raw (grey lines) and spatially corrected (blue lines) leaf area for the plants (replicates) of two genotypes, one per panel, under the two water regimes (as illustration). WD stands for water deficit and WW for well watered.

### Second stage results: PhenoArch platform

The three-level nested hierarchical growth model in equation (3) was fitted to the spatially corrected leaf area. We used cubic B-spline basis of dimension 13 to represent *f*_*p*_, *f*_*pg*_ and *f*_*pgi*_ (i.e., *b*_pop_ = *b*_gen_ = *b*_plant_ = 13). Under this configuration, the mixed model in equation (5) has 52992 observations (there are missing values), 23920 regression coefficients (both fixed and random) and 11 variance components. Estimation took approximately 9 minutes. Computations were performed in (64-bit) `R` 4.1.0, and a 2.40GHz × 4 Intel Core™ i7 processor computer with 15.6GB of RAM and Ubuntu 16.04 LTS OS.

Estimates at the three levels of the hierarchy were obtained: (1) population growth curves 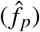, (2) genotype-specific deviations 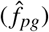 and respective growth curves 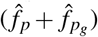, and (3) plant-specific deviations 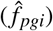 and respective growth curves 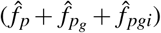. Note that the four populations (panel by water regime combination) show a different growth pattern. Plants in water deficit show a lower growth compared to well watered plants for both panels (Figure 4(a)). Genotype-specific deviations from their estimated population mean are shown in Figure 4(b) where positive and negative deviations refer, respectively, to better and worse genotypic performance compared to the mean population. As expected, the magnitude of the deviations (and, thus, the differences among genotypes performance) increases with time. Also, genotypes from Panel 1 show the largest genetic variation under both water regimes. This is in concordance with the spatially corrected data (grey lines in Figure 4(a)). The genotype-by-water regime interaction is illustrated for two genotypes per Panel in Figure 4(c). When compared to the average performance of the genotypes in the Panel, genotype 44 in Panel 1 and genotype 20 in Panel 2 behave similarly under both water regimes (they depict similar deviations under both treatments). In contrast, genotype 43 in Panel 1 presents a better performance under WD regime, while genotype 03 in Panel 2 displays a better performance under WW regime. Finally, the model is able to successfully recover the evolution over time of the spatially corrected leaf area (Figure 4(d), grey lines), while appropriately handling the missing data, as illustrated with the plant-specific growth curves (Figure 4(d), dotted blue lines) of two genotypes (see also results for the populations means in Figure 4(a)). Additionally, genotype-specific growth curves (Figure 4(d), orange lines) seem to properly summarise/describe the behaviour of the plant curves. To asses the relative accuracy of our model in predicting the plant-specific growth curves, i.e. 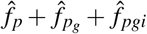, we computed the mean absolute percentage error (MAPE). For this data, the MAPE value was 2.9%, indicating a good performance.

**Figure 4.**
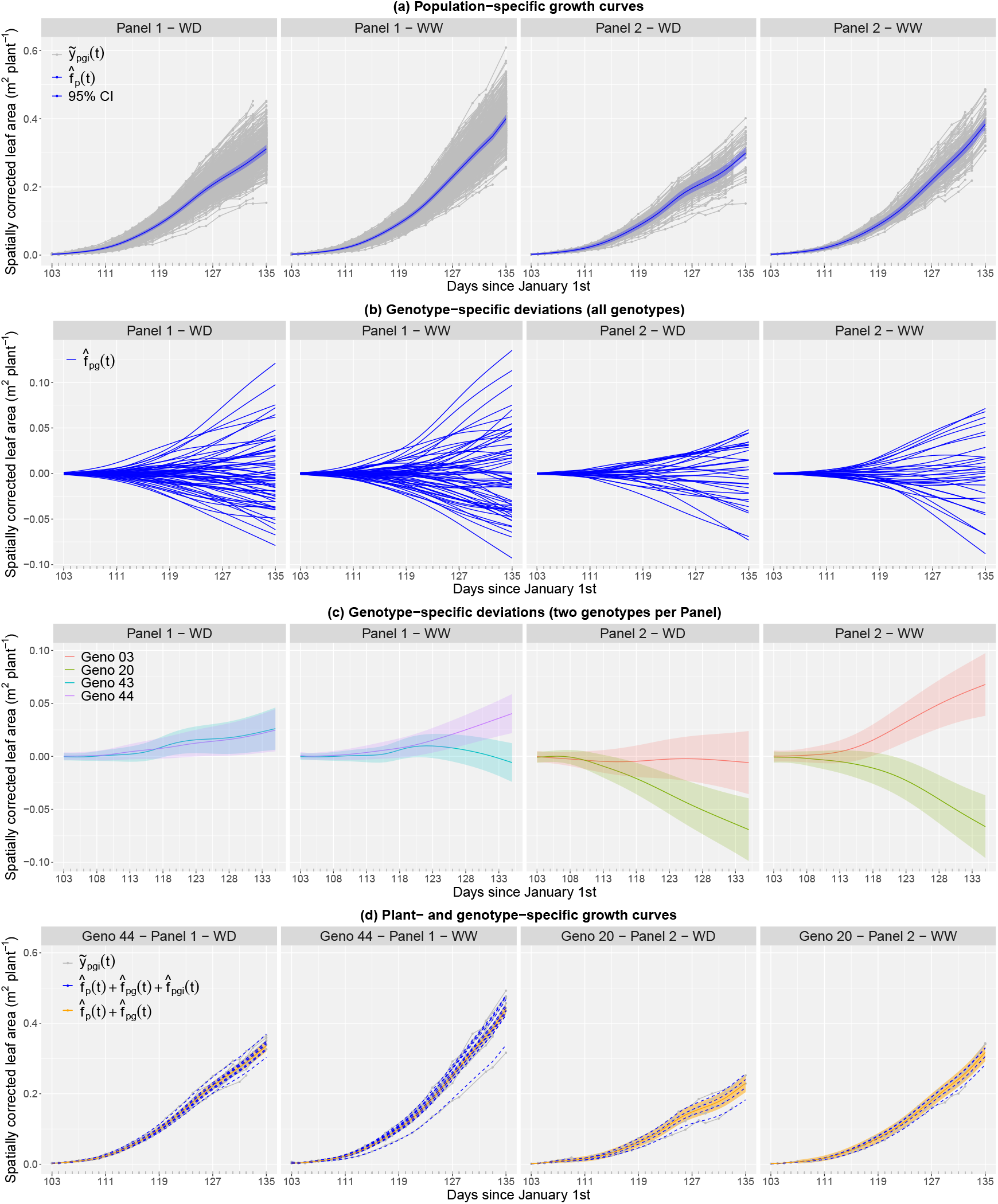
Results of the second stage of analysis for the PhenoArch platform: (a) Estimated population growth curves (continuous blue lines) with 95% pointwise confidence intervals (blue shaded areas), (b) estimated genotype-specific deviations for all genotypes, (c) estimated genotype-specific deviations for three genotypes per population (as illustration) with 95% pointwise confidence intervals (shaded areas); and (d) estimated plant-(dotted blue lines) and genotype- (orange continuous lines) specific growth curves with 95% pointwise confidence intervals (orange shaded areas) for two genotypes, one per panel, under the two water regimes (as illustration). WD stands for water deficit and WW for well watered. In (a) and (d) the grey lines represent the spatially corrected leaf area at the plant level (first stage).

#### FIP platform (ETH Zürich)

The FIP platform, located at the ETH research station in Lindau-Eschikon (Switzerland), is a cable-suspended multi-sensor platform designed for automated, accurate and supervised high throughput data acquisition on an area of 1 hectare^56^. From 2015 to 2017, the FIP platform was used to measure the development of canopy height on a diverse panel of European wheat genotypes (GABI wheat^57^), including a panel of Swiss varieties^15, 16, 51^. Here we focus on the 2017 data where height measurements started well before jointing. Figure 1(b) shows the FIP platform with its crop rotation allocated to six different lots; in 2017 the wheat experiment was planted in lots 2 and 6. Details on the experiment and experimental design can be found in Kronenberg *et al*. (2017, 2021)^16, 51^. In short, the experimental unit was a plot to which the genotypes were allocated as only treatment factor in an augmented 2D design. Three checks were placed in nine complete replications per lot, and test genotypes were allocated in a row-column design assuming that each row within a replicate (lot) received different environmental conditions (due to variability of crop husbandry measures, such as crop protection and fertilisation) while the upper, central and lower range of each lot received similar conditions (due to the similar slope direction within both lots). Canopy height measurements were carried out in irregular intervals of 2 to 13 days between February 27th and June 30th 2017 (which correspond to 58 and 181 DOY, respectively), using a terrestrial laser scanner mounted on the FIP sensor head^51^. To analyse the experiment, we arranged the two replicates (lots) diagonally in a virtual grid of *r* = 42 rows by *c* = 36 columns. Additionally, the country in which a genotype was first inscribed into the European variety catalogue was also considered in the analyses. We used this information to allocate the genotypes to different wheat populations targeted for specific regions within Europe. We will refer to these wheat populations (groups of genotypes) as regions. Accordingly, plots are nested in genotypes, and genotypes are nested in regions (wheat populations). The number of genotypes per region was as follows: Austria/Czechia (AT/CZ): 17; Switzerland (CH): 39; Germany (DE): 114; Sweden/Denmark (SE/DK): 11; France (FR): 90; Great Britain (GB): 38; Poland (PL): 25. The final dataset consists of *n* = 23 dates at which canopy height was measured on *M* = 720 plots (720 × 23 = 16560 observations, including missing data), *L* = 334 genotypes, and *k* = 7 populations (regions).

#### First stage results: FIP platform

In this case, the SpATS model included, besides the spatial trend and the random genotypic effects, fixed effects for the two lots (experimental design factor) and the seven wheat populations (region of origin), as well as random effects for the row and column positions. We also considered different genetic variances for each of the seven populations (regions). For the spatial trend, basis dimensions of 42 and 36 were assumed for the row and column positions of the virtual grid, respectively. The correction included the estimated population and genotypic effects and the residuals, and we averaged over the lot fixed effect to eliminate its impact. The spatially corrected canopy heights obtained when modelling genotypes either as fixed or random were, also here, very similar (see Supplementary Fig. S5 online). As for the PhenoArch experiment, the correction performed in the first stage reduced the variability among replicates of the same genotype (see Supplementary Fig. S4 and S6 online). This reduction is due to the lot effect and the spatial variation. The spatial distribution of the raw and spatially corrected canopy height, at four different measurement days (DOY), is shown in Figure 5.

**Figure 5.**
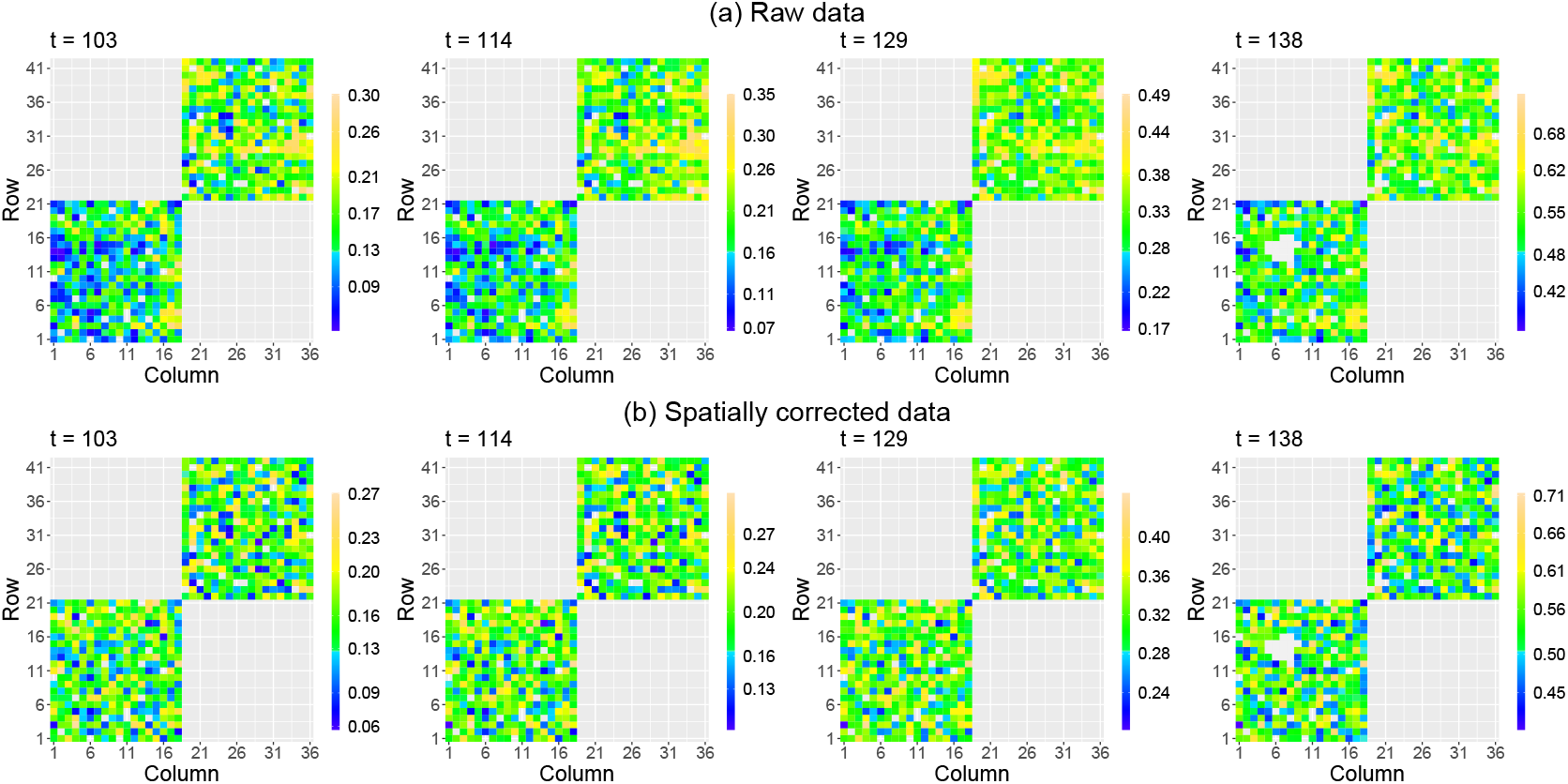
Results of the first stage of analysis for the ETH field phenotyping platform: Spatial distribution of the (a) raw and (b) spatially corrected canopy height at four different measurement days (*t* = 103, 114, 129, 138 DOY). The white areas denote missing data. The colour scale is different for each day.

#### Second stage results: FIP platform

For this analysis, and to allow the genetic variation to vary across regions (as suggested by the spatially corrected data; see Supplementary Fig. S4(b) online), we assumed, in equation (4), different values of 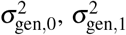 and 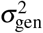 per region. The growth curves for the spatially corrected phenotype show here a more complex pattern than for the PhenoArch platform (see Supplementary Fig. S2(b) and S4(b) online), so we considered cubic B-spline bases of dimension 20 for the three levels of the hierarchy (*f*_*p*_, *f*_*pg*_ and *f*_*pgi*_). Under this configuration, mixed model in equation (5) has a total of 21220 coefficients, 16560 observations (including missing data) and 32 variance components. The fitting processes needed approximately 10 minutes. The MAPE was 1.94%.

In addition to the estimated growth curves and deviations at the three levels of the hierarchy, for this dataset we also obtained the first-order derivatives of the region and genotype-specific growth curves (Figure 6); they have shown to be good descriptors of genotype specific growth habit^52^ (recall that the first-order derivative is an indicator of the growth rate, i.e. the speed of canopy height development). Results show differences between regions (and between genotypes within a region) in, e.g., growth patterns (Figure 6(a)), growth rates (Figure 6(b)), and genotypes performance (Figure 6(c)). To better illustrate differences among regions, Figure 7(a) shows the estimated region growth curves together with the daily average temperature along the experiment. In Figure 7 (b) important maxima (i.e. largest growth rates) are indicated in the first-order derivatives of the respective region growth curves. Results for the replicate-specific growth curves are included in Supplementary Fig. S7 online (for one genotype per region, as illustration).

**Figure 6.**
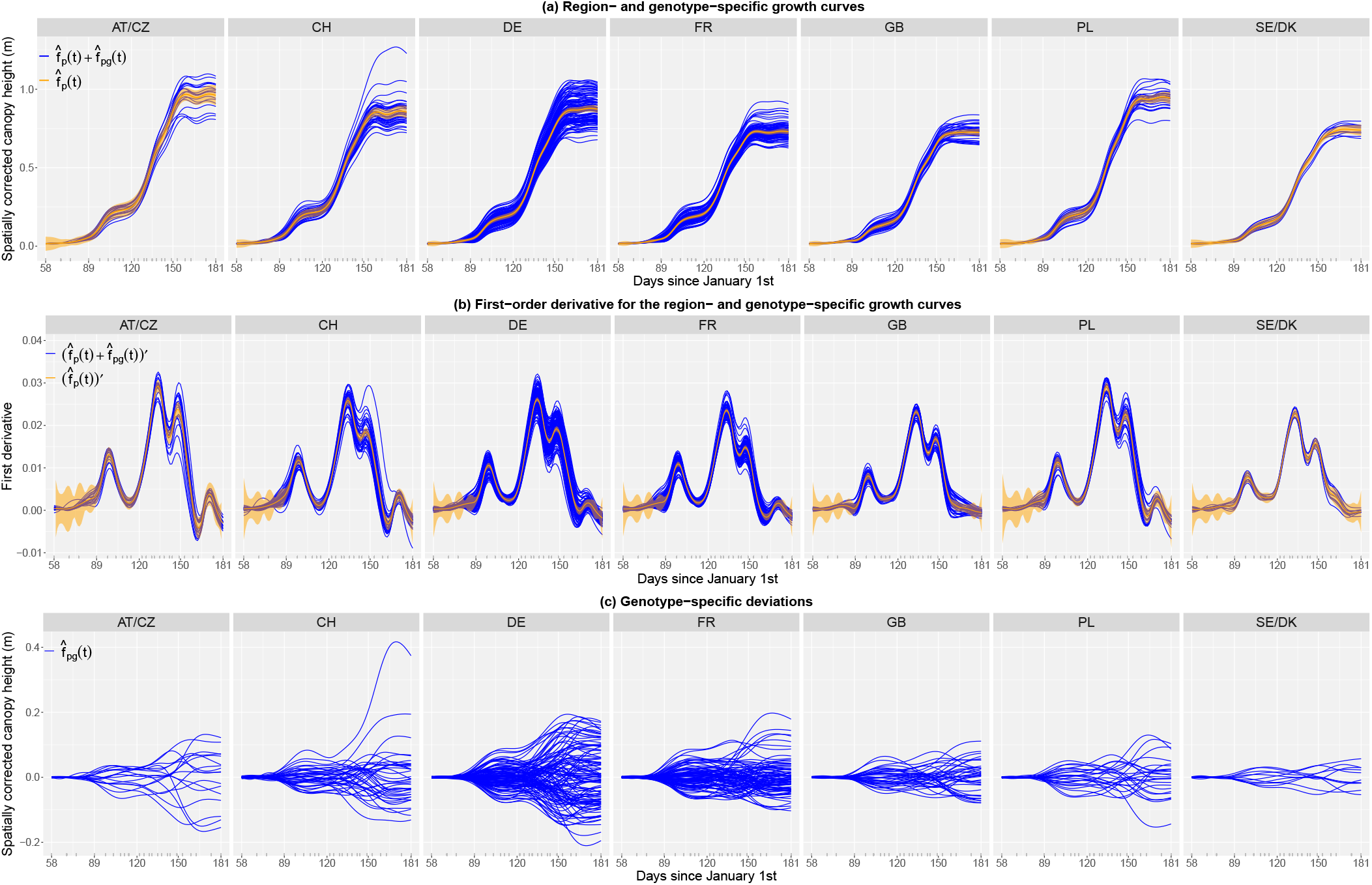
Results of the second stage of analysis for the ETH field phenotyping platform: (a) estimated region (orange) and genotype-specific (blue) growth curves, (b) estimated region (orange) and genotype-specific (blue) first-order derivatives, and (c) estimated genotype-specific deviations. In (a) and (b) the orange shaded areas denote 95% pointwise confidence intervals at the region level. AT/CZ: Austria/Czechia; CH: Switzerland; DE: Germany; FR: France; GB: Great Britain; PL: Poland; SE/DK: Sweden/Denmark.

**Figure 7.**
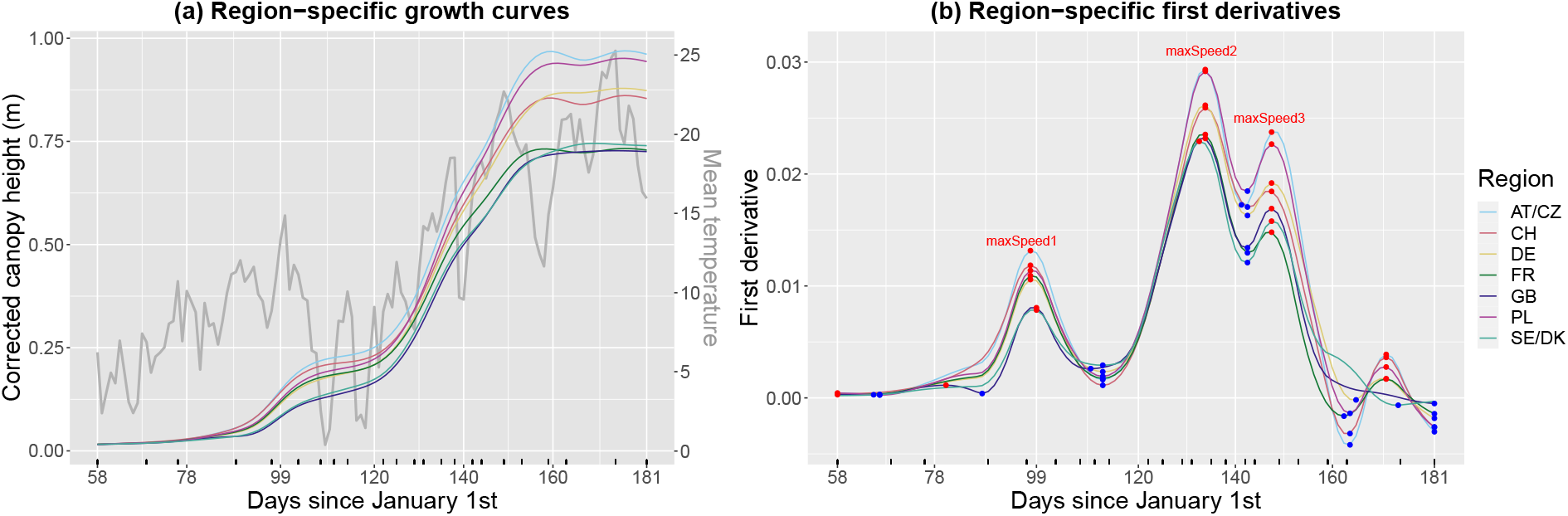
Results of the second stage for the ETH field phenotyping platform: (a) region-specific growth curves (coloured lines) vs. mean temperature (grey line), and (b) region-specific first-order derivatives; blue and red points indicate (local) minima and maxima, respectively. AT/CZ: Austria/Czechia; CH: Switzerland; DE: Germany; FR: France; GB: Great Britain; PL: Poland; SE/DK: Sweden/Denmark.

#### Extracting time-independent attributes to characterise genotypes

To characterise the genotypes, we extracted the maximum corrected canopy height (maxTrait) from the estimated genotype-specific growth curves (Figure 6(a)) as well as three maximum speed rates (maxSpeed1, maxSpeed2 and maxSpeed3) from their first-order derivatives (Figures 6(b) and 7(b)). These speed rates correspond to maxima around DOYs 97, 133 and 147, respectively. Maximum speed rates around DOY 133 (maxSpeed2) and 189 (maxSpeed3) can be interpreted, respectively, as recoveries after a severe cold period in April (DOY 110-120) and a milder one in May (DOY 140) (Figure 7(a), grey line). However, it appears that already for maxSpeed3 the growth rates declined as plants approached their final height. We also estimated the area under the genotype-specific deviations (Figure 6(c)) as a global measure of a genotype performance over time when compared to the genotypes of the same region. A positive (negative) area indicates a genotype performance better (worse) than the regional average. Here, the area (AUC) was estimated for the complete time interval where the genotypes were measured. Nothing, however, precludes to focus attention on a restricted time interval of interest.

The bivariate scatterplots of the extracted genotype-specific attributes show that the genotypes cluster according to their region of origin (Figure 8, lower off diagonal, one color per region). Some atypical genotypes can also be identified, for example an old, tall (>1.2m) Swiss variety. In the upper off diagonal, we report the Pearson’s correlation coefficient (marginal and by region). The strongest marginal correlations are between maxSpeed3 and maxTrait, and maxSpeed2 and maxTrait; maxSpeed3 and maxSpeed2 also present a high correlation. It is noteworthy that the marginal correlation between maxTrait and AUC is lower than the conditional correlation by region. Univariate analysis of each attribute using boxplots also shows regional clustering, and that clusters might change along time (e.g., compare clusters in maxSpeed1, maxSpeed2 and maxSpeed3).

**Figure 8.**
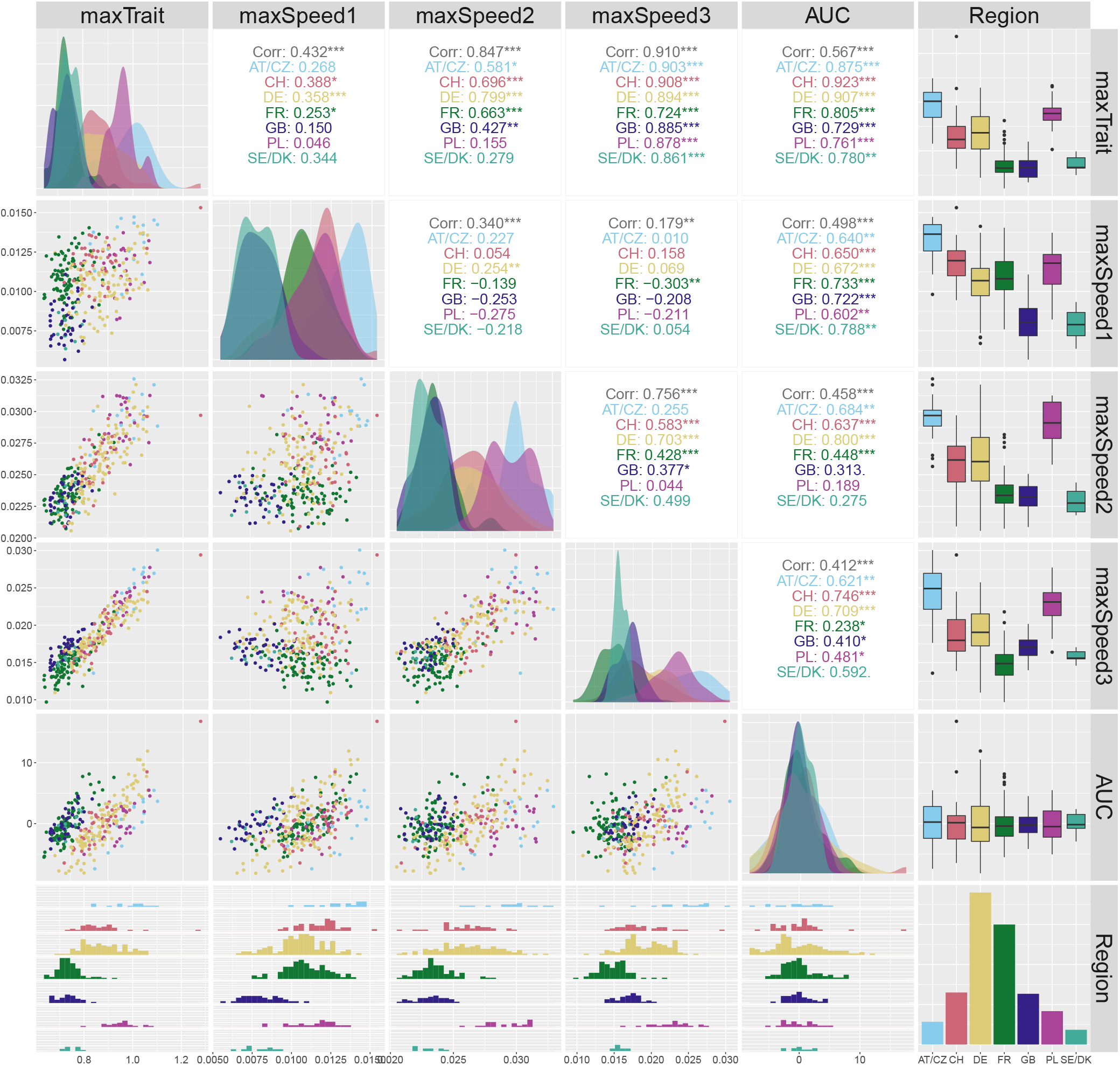
Scatterplots matrix with the extracted attributes at the genotype level for the ETH field phenotyping platform. The lower off diagonal depicts bivariate scatterplots, the diagonal shows the conditional densities of each attribute per region, the upper off diagonal indicates the bivariate Pearson correlation (marginal and by region), the last column displays the boxplots of each attribute per region, the last row depicts the conditional histograms of each attribute per region, and the bottom right barplot shows the number of genotypes per region. AT/CZ: Austria/Czechia; CH: Switzerland; DE: Germany; FR: France; GB: Great Britain; PL: Poland; SE/DK: Sweden/Denmark.

#### Use of time-independent attributes to characterise regional adaptation

While a deeper physiological analysis is beyond the scope of this publication, we will use the extracted attributes to highlight the potential benefit of an in-depth analysis of spline-based growth patterns. Here, we use the regional groups but a similar analysis could be done using individual genotypes. The observed height development follows a principally logistic growth curve: stem elongation started after the plants were vernalised over winter (by means of cold exposure) and ended around flowering. However, the height development plateaued between day 103 and 118 in all genotypes, most likely due to a cold period in April. When looking at the region specific growth curves, this short and extreme phase caused even rank changes in growth (first-order derivative in Figure 7): the regional groups showing most vigorous growth before the stress (maxSpeed1) stopped growth completely while those which grew slowest could maintain some growth during the cold (first local minima following maxSpeed1). Such pattern may point to physiological adaptations to the different climatic regions of Europe as the slow-growing northern types from Great Britain (GB) and Denmark and Sweden (SE/DK) showed least response to cold while the fast-growing continual types from Poland (PL) and Austria and Czechia (AT/CZ) stopped growing. Moreover, the genotypes from the south-west – France (FR) and Switzerland (CH) – did not recover growth up to the same level as the more northern and eastern varieties did (compare maxSpeed1 with maxSpeed2). We acknowledge that a solid interpretation of this pattern and its significance will require a multi-year analysis to sample the typical genotype or region-specific average development.

## Discussion

In agricultural and breeding research, non-destructive data acquisition of phenotypic traits by HTP platforms has emerged in recent years as a rich source of new information on plant growth and development as well as on genotypic performance. This has been accompanied by the need for novel and appropriate statistical methods of analysis. In this paper, we focus on the analysis of a common type of HTP data in the form of time-series observed on phenotypic traits that are still close to the platform measurements (low-level traits), with these time-series occurring within a hierarchical structure, of plants or plots nested in genotypes, and genotypes nested in populations (of genotypes). We propose a statistically flexible and computationally feasible method that decomposes the required spatio-temporal analysis into two stages. In the first stage, we correct the “raw” HTP data for (nuisance) spatial variation and obtain spatially corrected time-series at the resolution of plants or plots with reduced between replicates/plots variability. The second stage consists of a temporal analysis with a hierarchical curve data model to jointly estimate growth curves at each level of the hierarchy (plant or plot, genotype, and population) as well as their first-order derivatives. From the estimated curves and their derivatives, we calculate new phenotypic traits, attributes, that we call intermediate traits. Although we have not explicitly covered it in the paper, these new phenotypic traits, as well as the estimated curves, can be used for selection purposes in plant breeding, i.e., to differentiate between genotypes. For instance, estimates for intermediate-level traits can be used as genotypic covariates in models for higher-order phenotypic traits as described by van Eeuwijk *et al*. (2019)^2^. Higher order phenotypic traits can be understood as functions of low and intermediate traits, for either or both of biological and statistical reasons. For example, yield can be interpreted as a high-order phenotypic trait that can be modelled as a function of yield components, where the yield components may represent low and intermediate level phenotypic traits. In Roth *et al*. (2021)^34^, intermediate traits obtained from modelling HTP data are included into genotype-by-environment interaction analyses, and Moreira *et al*. (2020)^58^ discuss the use of information obtained from HTP time-series traits for genomic selection and the detection of QTL and causal variants. Finally, while we have focused in the paper on data with a nested structure, the proposed modelling framework can easily be extended to accommodate more complex structures, such as data with crossed levels of grouping^21^ (e.g., when modelling genotype-by-treatment interactions is of interest).

Regarding the statistical methods used at each stage of our modelling approach, the first stage relies on the SpATS model, and (hierarchical) P-splines are used for the second stage. However, our approach is flexible with respect to the choice for both the first and second stages. For instance, the separable autoregressive model^6^ represents a clear alternative for the first stage, while, for the second stage, hierarchical functional principal component analysis can be used^28^. We believe that our double P-spline approach is attractive both computationally and for interpretation, and the HTP data that we analysed in this paper and in other projects show that it works well (see Millet *et al*., 2021^20^ and https://eppn2020.plant-phenotyping.eu/ for more examples). Nevertheless, fairness requires us to mention some limitations as well. The P-spline hierarchical data model used in the second stage relies on specifying each function in the model using B-spline basis expansions. The model contains, consequently, a large number of parameters. Typically, computational times are within an acceptable range (for the experiments analysed in this paper, around 10 minutes). Yet, the approach may not scale well to experiments where the number of plants (and associated basis dimension) is very large (due to the size of the system of equations to be solved). Regarding the number of B-spline basis functions, we recommend using the same value for the three levels of the hierarchy, even if this increases computation. The final number does not seem to impact results (estimated curves) significantly, provided it is large enough to capture the underlying patterns. However, the estimated first-order derivatives have shown to be more sensitive to the number of basis functions.

In the first stage of our two-stage approach, analyses are done separately per time point. As such, information on spatial heterogeneity is not shared across different measurement times, and there is the need that all plants or plots are measured at (approximately) the same times. From a modelling perspective, the simultaneous modelling of spatial and temporal genetic and non-genetic variation in a one-stage model will serve to share information on common spatial variability across measurement times and it may solve the problem of variable measurement times. Yet, one-stage approaches have the limitation of being very computationally demanding, especially when the number of observations and/or the parameters to be estimated is very large. Preliminary and proof-of-concept results can be found in Pérez *et al*. (2020)^14^ and in Verbyla *et al*. (2021)^13^. In a similar vein to the work presented here, Pérez *et al*. (2020)’s approach is based on using three-dimensional P-splines for modelling the non-genetic spatio-temporal (noise) process and hierarchical P-splines for the longitudinal genetic signal. In Verbyla *et al*. (2021), multi-dimensional splines are also used for the spatio-temporal noise, whereas a factor analytic structure is considered for the genetic effects. Although promising, both approaches suffer from severe computational limitations (e.g., Verbyla *et al*. 2021, report convergence requiring several days). More work is thus required to render them feasible for routine use, and to date, stage-wise approaches represent not only computationally feasible but also statistically valid alternatives.

Apart from this work, van Eeuwijk *et al*. (2019)^2^ also proposed a two-stage approach for the analysis of HTP data, where they first correct for spatial variation, and then focus on estimating, and further processing, the temporal dynamic of the genetic effects. In that paper, spatially adjusted genotypic means are carried to the temporal analysis. In contrast, our proposal allows keeping the data resolution for the second stage at the experimental unit. Also, in the second stage, we jointly model the whole sample of spatially corrected growth curves, while in van Eeuwijk *et al*. (2019)^2^ analyses are done separately per genotype. Our hierarchical approach thus allows borrowing strength across plant/plot curves for more efficient estimation of genotype and population growth trajectories. This is particularly important in the presence of incomplete data. We are aware that when choosing a two-stage approach, one might also choose, as done in Roth *et al*. (2021)^34^, to first model the longitudinal variation at a plant or plot level and subsequently apply a spatial correction to extracted features. In that paper, a P-spline model is first fitted separately for each plot time-series, from which the timing of key stages (among other features) are extracted. These intermediate traits are then processed to obtain spatially adjusted genotypic means for further analyses. We feel that both options - with the spatial or the temporal analysis first - represent valid alternatives. The choice for one or another method will depend on the relative magnitudes of the various spatial, temporal and spatio-temporal genetic and non-genetic processes and may be difficult to assess beforehand. A study of the proposal that best suits particular situations represents an interesting area of study. It should be realised that stage-wise analyses make it necessary to propagate uncertainty from stage to stage. Here, we did it by weighting the second stage with the (inverse) of the estimated variance associated with the spatially corrected trait. In the HTP context, weighting has shown to improve results^34^.

All in all, the two-stage approach described in this paper represents a good compromise between accuracy, adequacy, computational efficiency and interpretability. The results show that our proposal is feasible on standard computers and delivers good descriptions of the genetic and non-genetic variation in the temporal dimension together with useful summary statistics for breeding purposes. We believe that it represents a powerful tool for routine application in phenotyping experiments with dense time series. To allow practitioners to use our proposal, the `R`-functions implementing it are publicly available at https://gitlab.bcamath.org/dperez/htp_two_stage_approach, where we also provide the code to reproduce the analyses and results shown in the paper.

### Three-level Nested Hierarchical Growth Model: Mixed Model Formulation

On the basis of model in equation (3) and specification in equation (4), we use the notation 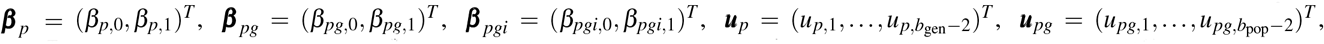, 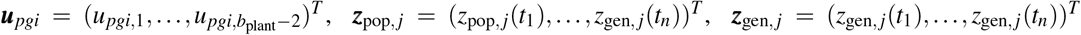 and ***z***_plant, *j*_ = (*z*_plant, *j*_(*t*_1_), …, *z*_plant, *j*_(*t*_*n*_))^*T*^. Then, in matrix notation, the three-level nested hierarchical growth model can be expressed as

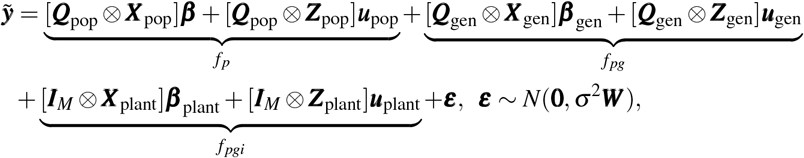

where ⊗ is the Kronecker product, ***Q***_pop_ and ***Q***_gen_ are matrices assigning, respectively, plants to populations and plants to genotypes, and

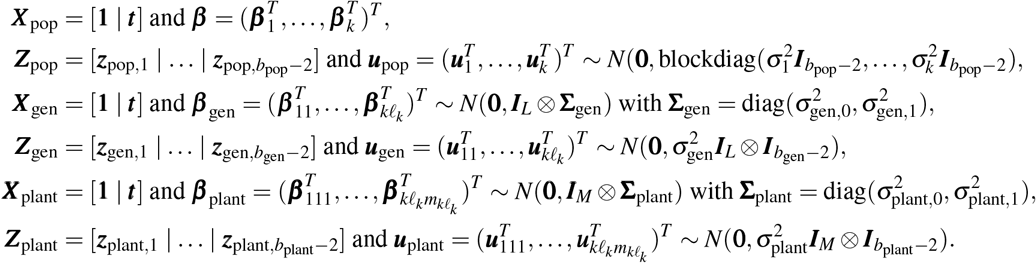

Finally, ***W*** is diagonal matrix whose diagonal entries are the weights from the first stage. We finish by noting that, under the previous model specification, there is the assumption that observations arising from the same plant, genotype or population are serially correlated, and the correlation increases as a function of the shared grouping levels^21^. In particular, curves 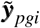 and 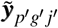 have covariance

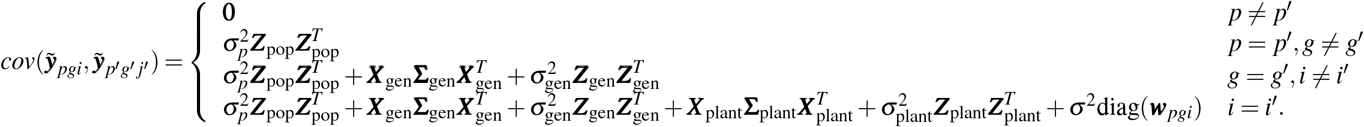

## Data availability

Both the PhenoArch and FIP datasets analysed during the current study are available within the article and its “Supplementary information”.

## Code availability

The `R`-functions implementing the two-stage approach, as well as the `R`-codes needed to reproduce the analyses for the PhenoArch and FIP platforms are available within the article and its “Supplementary information”.

## Author contributions statement

D.M.P.: methodology, software, data curation, data analysis, visualization, original draft, review and editing. M.X.R.: conceptualization, methodology, software, review and editing, supervision. M.B.: methodology, software, review and editing. A.H: data collection, review and editing. L.K.: data collection, data curation, review and editing. Ll.C.B.: data collection, data curation, review and editing. E.M.: methodology, software, data curation, review and editing. F.E: conceptualization, methodology, review and editing, supervision. All authors reviewed and approved the manuscript.

## Supplementary information

Supplementary information (code and data) is available for this paper in the Basque Center for Applied Mathematics repository, https://gitlab.bcamath.org/dperez/htp_two_stage_approach.

