## Supplementary figures for "A two-stage approach for the spatio-temporal analysis of high-throughput phenotyping data"

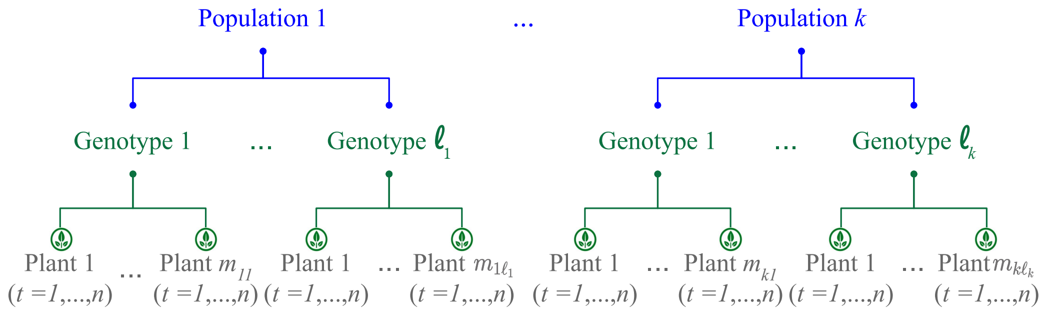


**Supplementary Figure S1.** Three-level nested hierarchical data structure

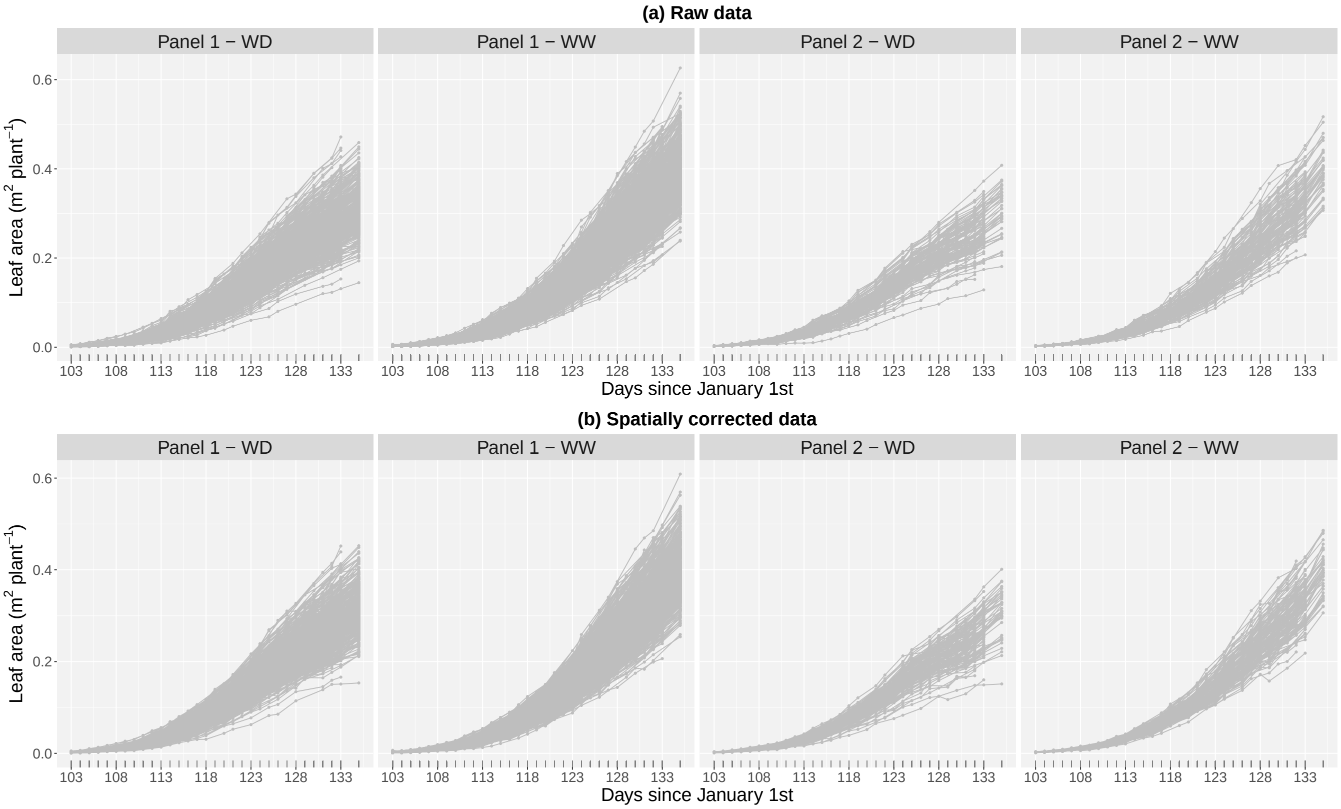


**Supplementary Figure S2.** For the PhenoArch platform: Evolution over time of the (a) raw leaf area and (b) spatially corrected leaf area. Results are shown for all plants in the experiment.

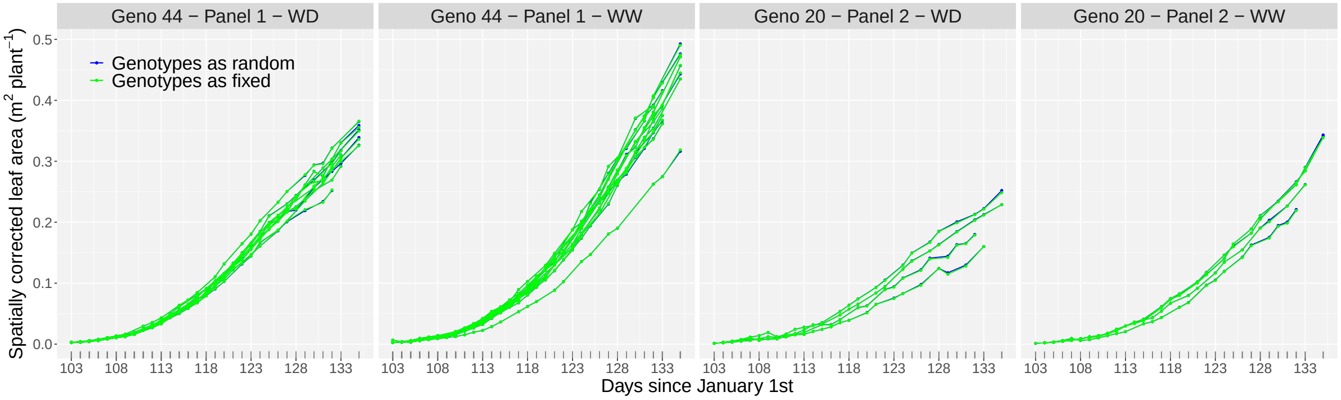


**Supplementary Figure S3.** For the PhenoArch platform: Comparison of the evolution over time of the spatially corrected leaf area when modelling, in the first stage, genotypes as random (blue lines) or fixed (green lines) effects. Results are shown for the plants of two genotypes, one per panel, under the two water regimes (as illustration).

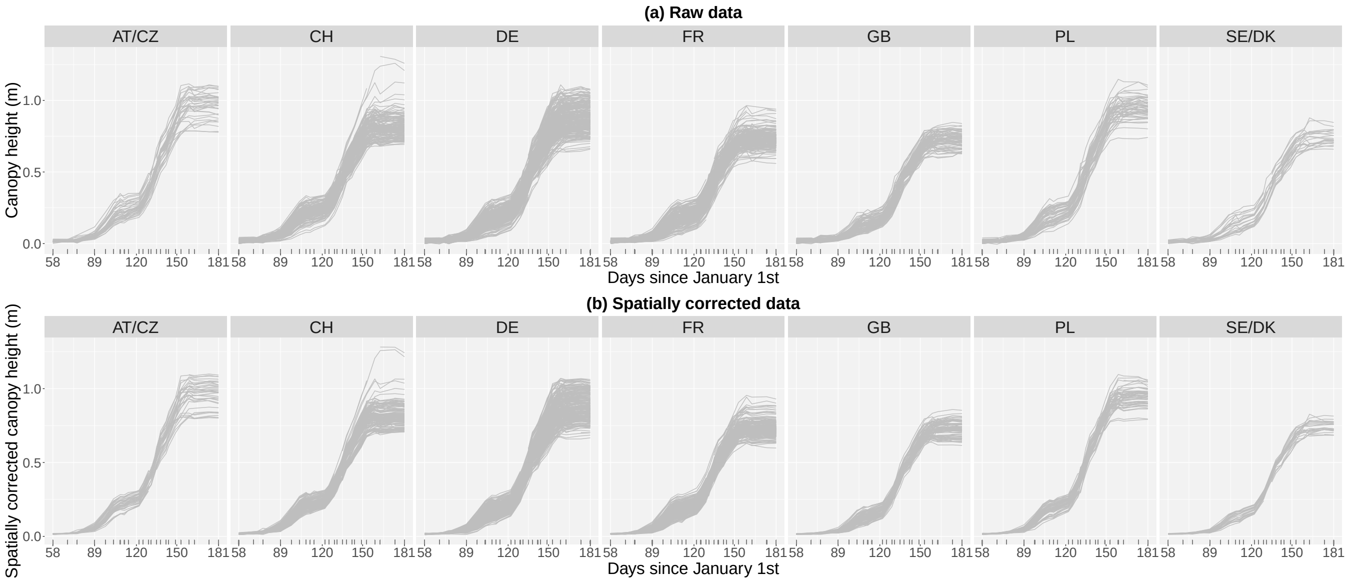


**Supplementary Figure S4.** For the ETH field phenotyping platform: Evolution over time of the (a) raw canopy height and (b) spatially corrected canopy height. Results are shown for all plots in the experiment.

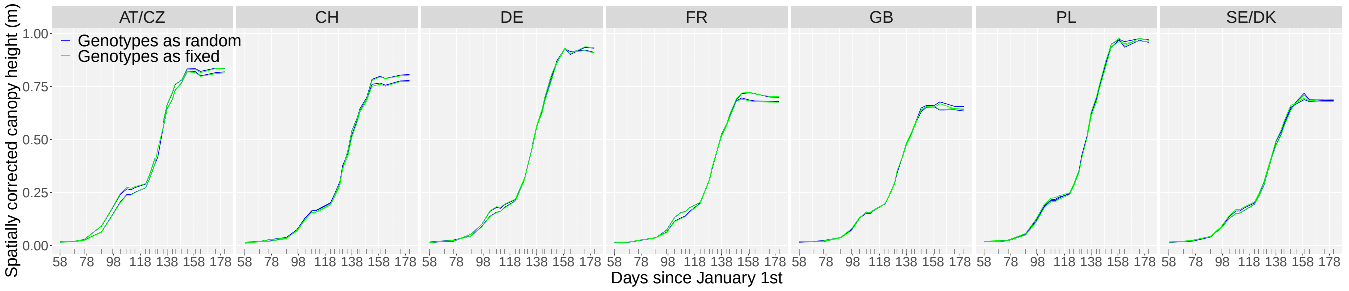


**Supplementary Figure S5.** For the ETH field phenotyping platform: Comparison of the evolution over time of the spatially corrected canopy height when modelling, in the first stage, genotypes as random (blue lines) or fixed (green lines) effects. Results are shown for the replicates of seven genotypes, one per region (as illustration).

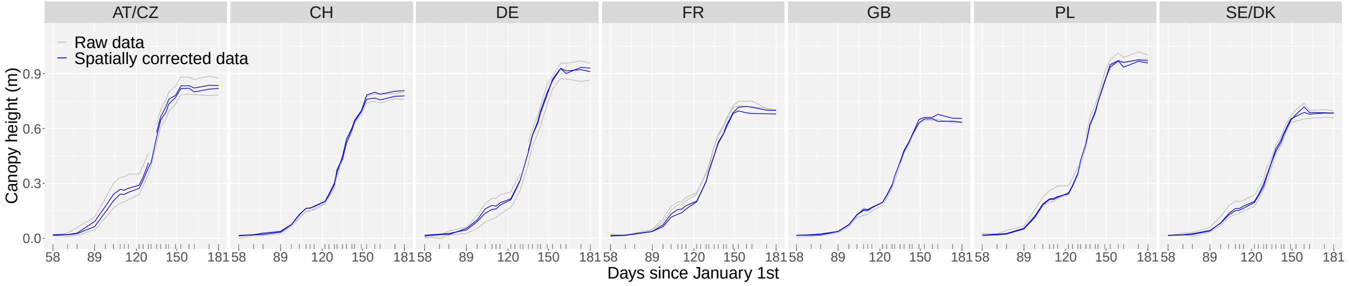


**Supplementary Figure S6.** Results of the first stage for the ETH field phenotyping platform: Evolution over time of the raw (grey lines) and spatially corrected (blue lines) canopy height for the replicates of seven genotypes, one per region (as illustration).

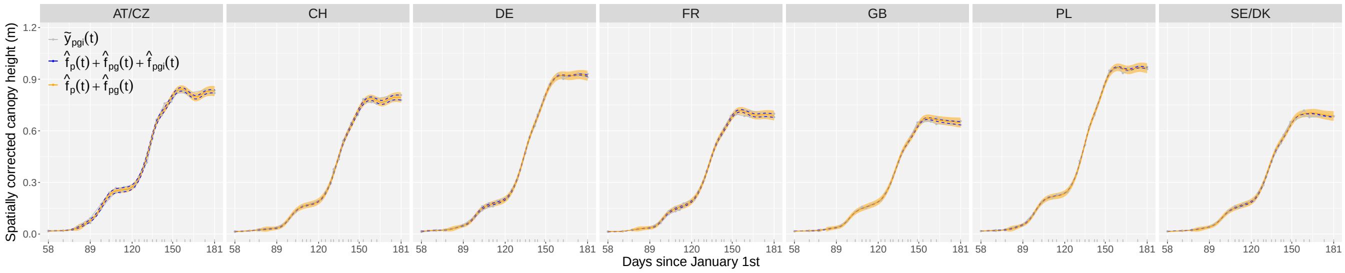


**Supplementary Figure S7.** Results of the second stage for the ETH field phenotyping platform: Estimated replicate- (dotted blue lines) and genotype- (orange lines) specific growth curves for one genotype per region (as illustration). For the genotype-specific growth curves, the orange shaded areas represent 95% confidence intervals. The grey lines represent the spatially corrected canopy height at the replicate/plot level.
